# Lineage-specific expansion of a potential novel class of resistance genes in Myrtaceae

**DOI:** 10.64898/2026.07.23.740426

**Authors:** Tamene T Tolessa, Scott Ferguson, Ashley Jones, Xiaoxiao Zhang, Simon Williams, Zhenyan Luo, Peri A Tobias, Shubiao Wu, Justin O. Borevitz, Benjamin Schwessinger, Rose Andrew

**Affiliations:** School of Environment and Rural Science, University of New England, Armidale, New South Wales, Australia; Research School of Biology, Australian National University, Canberra, Australian Capital Territory, Australia; Department of Integrative Biology, University of California, Berkeley, CA, USA; School of Life and Environmental Sciences, University of Sydney, Camperdown, NSW 2006, Australia

**Keywords:** Myrtaceae, NLR, resistance genes, non-canonical domains, Jacalin

## Abstract

Genes encoding intracellular nucleotide-binding site leucine-rich repeat (NBS-LRR) receptors represent the largest class of resistance (*R*) genes in plants, yet their evolutionary trajectory in trees remains poorly understood. Using high-quality long-read genome assemblies from eight Myrtaceae species, we identified 15,792 NBS-encoding genes with threefold variation in gene content across species.

To investigate potential decoy domains for effector proteins of pathogens, we parsed the annotated NBS-encoding genes and determined 181 unique non-canonical domains. Notably, we observed frequent integration of Jacalin domains into TIR-NBS proteins. This novel gene family, named TNJ, is hypothesized to represent a new *R* gene class.

Further exploration of TNJ sequence analysis shows up to seven Jacalin domains per protein and forming a monophyletic clade; however, AF3 modelling confirmed only six domains. Conserved residues in the TIR domain and functional motifs within the NB-ARC domain supports TNJ’s potential role in immune signalling.

Most hypervariable sites and positively selected sites detected to be surface-exposed were clustered in the Jacalin region of TNJ, suggesting that surfaces of Jacalin domain may harbour residues determining pathogen recognition specificity. These findings support TNJ as a potentially new class of *R* genes in Myrtaceae with Jacalin as a replacement for LRR.

## Introduction

Plant disease resistance (*R*) genes encode receptor proteins with a modular architecture characterized by a conserved nucleotide-binding sites (NBS) and the C-terminal leucine-rich repeats (LRR) (Van de Weyer *et al*., 2019; Kourelis *et al*., 2021), hereafter called *NLRs*. These genes play essential roles in plant defence against diverse disease-causing pathogen effector proteins, including those from bacteria, fungi, oomycetes, nematodes and viruses (van der Hoorn and Kamoun, 2008; Kourelis and van der Hoorn, 2018; Araújo *et al*., 2019). Based on variations in their N-terminal domains, NLRs are classified into TIR-NBS-LRR (*TNL*), CC-NBS-LRR (*CNL*), and RPW8-NBS-LRR (*RNL*) (Meyers *et al*., 2003; Van de Weyer *et al*., 2019), many of which contain additional functional domains or motifs (Kroj *et al*., 2016; Sarris *et al*., 2016; Bailey *et al*., 2018; Zhang *et al*., 2022). The modular structure of NLRs facilitates domain swapping (Cesari *et al*., 2014; Kroj *et al*., 2016; Marchal *et al*., 2022), duplication (Shao *et al*., 2016; Dolatabadian *et al*., 2022), and integration of novel domains (Bailey *et al*., 2018; Van de Weyer *et al*., 2019; Marchal *et al*., 2022), driving NLR diversification (Dolatabadian *et al*., 2020; Chia *et al*., 2024) and new recognition specificities (Marchal *et al*., 2018; De la Concepcion *et al*., 2021; Cesari *et al*., 2022). A functional NLR protein contains conserved motifs critical for immune function, including P-loop, Kinase-2, RNBS-A to D, and GLPL motifs in the NBS domain, which are essential for ATP binding (Romero Romero *et al*., 2018; Wang *et al*., 2019). The amino acid residues required for pathogen recognition and enzymatic activity underpin the molecular mechanisms by which functional NLR proteins trigger hypersensitive cell death and confer disease resistance. Conserved enzymatically important residues are responsible for producing signalling molecules involved in plant defence (Huang *et al*., 2022; Jia *et al*., 2022; Lapin *et al*., 2022; Yu *et al*., 2022), and solvent-exposed variable residues that undergo rapid positive selection are crucial for pathogen effector recognition (McHale *et al*., 2006; Prigozhin and Krasileva, 2021).

Non-canonical domain (NCD) integration into NBS-containing proteins is widespread across monocots, dicots, and mosses, often occurring at N- or C-terminal or inter-domain positions, suggesting an important role of NCDs in NLR diversification and evolution, though their frequencies vary among species (Cesari *et al*., 2014; Kroj *et al*., 2016; Sarris *et al*., 2016; Bailey *et al*., 2018). Recent genome-wide comparative studies have found NCDs integration to have greatly expanded and diversified NLR repertoires, as seen in *Capsicum annuum* and *Brassica napus* (Kim *et al*., 2017; Ning *et al*., 2024). Several NCDs identified in NBS-containing proteins of flowering plants are proposed to act as baits for pathogen effector recognition (Kroj *et al*., 2016; Sarris *et al*., 2016; Li *et al*., 2019), though many remain functionally uncharacterized. In *Oryza sativa*, for instance, over expression of the jacalin-related Mannose-Binding Lectin (OsMBL) protein enhanced resistance to *Magnaporthe oryzae* by activation of chitin-induced reactive oxygen species burst and rice defence-responsive genes (Han *et al*., 2018). In Solanaceae, an extended N-terminal domain integrated into CNLs in a clade-specific manner (Seong *et al*., 2020) mediates effector interaction and oligomerisation (Li *et al*., 2019). Likewise, five C-terminal jelly roll/Ig-like domains (C-JID) encoded in the *Ma*, a *TNL* gene from *Prunus cerasifera*, confer complete resistance to root-knot nematode (Maruta *et al*., 2022). A recent pangenomic evolutionary analysis in *Sorghum bicolor* revealed that RMES1 proteins lacking N-terminal domains exhibit high polymorphism at the NBS P-loop motif. These proteins are thought to possess an NCD that activates the immune response network to counter a phloem feeding aphid, contributing to evolutionary rescue of *S. bicolor* (VanGessel *et al*., 2025). Although NCD integration during plant evolution has driven extensive NLR diversification, creating novel domain architectures with enhanced pathogen recognition capacity (Duxbury *et al*., 2016; Kroj *et al*., 2016; Bailey *et al*., 2018; Van de Weyer *et al*., 2019; De la Concepcion *et al*., 2021; Maidment *et al*., 2023; Reveguk *et al*., 2025), the detailed diversity of NBS-encoding genes and their NCDs remain poorly explored in non-model plant lineages, particularly in the tree species.

Forest trees in both natural ecosystems and plantations are exposed to a wide range of biotic and abiotic stresses that threaten their survival and productivity. In recent decades, the rate of introduction and spread of invasive pests and pathogens have intensified with devastating consequences (Teshome *et al*., 2020). For instance, *Cryphonectria parasitica*, the causal agent of chestnut blight, has nearly eradicated North American chestnuts, (Newhouse *et al*., 2014), while *Fusarium circinatum* has caused severe pitch canker outbreaks in pine plantations worldwide (Gordon and Reynolds, 2017). Climate change induced weather extremes further exacerbate these threats, promoting widespread pest and disease epidemics, as exemplified by the destructive mountain pine beetle (Teshome *et al*., 2020; Fensham and Radford-Smith, 2021). More recently, the fungal pathogen *Austropuccinia psidii* (myrtle rust) has emerged as a major global threat to the Myrtaceae family, a diverse lineage comprising around 5,800 species of high ecological and economic importance across tropical and subtropical regions (Roux *et al*., 2016; Saber *et al*., 2024). The increasing challenges highlight the urgent need for proactive mitigation strategies, including advancing genomic knowledge of *NLR* genes to support breeding programs that enhance sustainable disease and pest management in forest species.

Around 450 such functionally validated *NLRs* have been identified in crops and model plants (Chia *et al*., 2024). An in-depth genome-wide analysis revealed distinct *R* gene classes, including NL-only domains in Arabidopsis (Van de Weyer *et al*., 2019), an autonomous ancient *NLR* (ANLs) with auto-active coiled-coil (CC) domain in *Capsicum annuum* (Lee *et al*., 2020), and a Zf-BED-NBS-encoding novel resistance profile forming distinct phylogenetic clade in *Populus trichocarpa* (Yang *et al*., 2008). Despite advances in genome sequencing and the growing availability of genome data (Ferguson *et al*., 2024), *NLR* gene families remain largely uncharacterized in the Myrtaceae family including in the iconic *Eucalyptus* species. Although some resistance QTLs (Junghans *et al*., 2003; Mamani *et al*., 2010; Thumma *et al*., 2013), and NLRs have been identified and classified in Myrtaceae, including within *Eucalyptus grandis* (Christie *et al*., 2016), *Syzygium maire (Balkwill et al., 2024)*, *Syzygium luehmannii (Tobias et al., 2025), Melaleuca quinquenervia* (Chen *et al*., 2023), and *Melaleuca alternifolia* (Chakrabarty *et al*., 2023), the broader diversity and evolutionary dynamics of NBS-encoding genes and their NCDs remain largely unexplored.

This study aims to investigate the evolutionary and functional implications of the novel domain structure containing the NCD Jacalin at the C-terminal of TIR and NBS (TN) proteins, which we refer to as TNJ. We performed a genome-wide analysis of NBS-encoding proteins across eight long-read sequenced species representing four Myrtaceae tribes (Thornhill *et al*., 2015) to characterise domain structure in TNJ. Phylogenetic analysis was used to address the evolutionary origin of TNJ. We then asked whether TNJ had retained the capacity to function as an R protein by identifying conserved functional elements (Romero Romero *et al*., 2018; Wang *et al*., 2019; Huang *et al*., 2022; Jia *et al*., 2022; Lapin *et al*., 2022; Yu *et al*., 2022) in the TIR and NBS domains of TNJ. Finally, we asked whether the Jacalin domains contained hypervariable sites and signatures of positive selection that could contribute to receptor specificity. Our study presents an important mechanism of lineage-specific expansion of new plant *R*-gene family by tandem duplication of Jacalin domain and enhances our understanding of the dynamic plant NLR diversity and evolution.

## Materials and Methods

### Annotation of NLR and non-canonical domains (NCDs)

We used long-read-sequence based genome assemblies of eight species: *Eucalyptus albens, Eucalyptus grandis, Angophora floribunda, Psidium guajava, Rhodaminia argentea, Rhodomyrtus psidioides, Syzygium oleosum and Melaleuca quinquenervia*. These species are representing four well-supported tribes of Eucalypteae, Myrteae, Zyzygieae and Melaleuceae (Table **S1**). Assembly completeness of *E. albens, E. grandis* and *A. floribunda* (Eucalypteae species) were reported in (Ferguson *et al*., 2024). The remaining five genomes were separately assessed using BUSCO (Table **S2**). To annotate predicted NBS-encoding genes, we employed a comprehensive NBS-encoding protein annotation pipeline, FindPlantNLRs (Chen et al., 2023) on unmasked genomes. The pipeline first uses NLR-ANNOTATOR (Steuernagel *et al*., 2020) to capture genomic regions of *NLR*-loci including their coordinates. It then implements homology searches of a tBLASTn translated nucleotide database using 235 experimentally validated NLR amino acid sequences (Kourelis *et al*., 2021) and uses ‘bedtools getfasta’ to obtain nucleotide sequences of the identified *NLR* loci for gene model prediction.

To annotate gene models from the identified *NLR*-loci, the FindPlantNLRs runs BRAKER2 (Hoff *et al*., 2019) in GeneMark-EP (Brůna *et al*., 2021) mode. To train gene model prediction, we used 3,842 protein hint sequences from orthogroups of *NLR* genes generated from 16 different plant species using OrthoFinder version 2.5.4 (Emms and Kelly, 2019) and protein sequences of 152 experimentally confirmed reference *R* genes (Osuna-Cruz *et al*., 2017). To search for the conserved NB-ARC domains against the predicted protein coding sequences of the gene model, the pipeline runs HMM search (Eddy, 2011) and screens the identified NB-ARC associated amino acid sequences for all NLR protein domains using InterProScan (Jones *et al*., 2014).

We used a separate custom python script to categorize different classes of NBS-encoding genes. The script parses the CATH-Gene3D domain structure database v4.2.0 (Buchan *et al*., 2002), Pfam v33.1 (Finn *et al*., 2014) and Coils (Lupas *et al*., 1991) domain identifiers from the output of InterProScan v-5.46-81.0 (Jones *et al*., 2014) following the description in Kourelis *et al*. (2021).

To identify the non-canonical domains (NCDs) in the predicted NBS proteins of each species, we parsed the PfamScan outputs corresponding NBS-encoding genes using the scripts ‘K-parse_Pfam_dmains_v3.1.pl’ and ‘K-parse_Pfam_domains_NLR-fusions-v2.4.2.pl’ (Sarris et al., 2016). Summaries and associated plots were created using R Statistical Software v4.3.0 (R Core Team, 2020).

### Analysis of TNJ domain structure, functional motifs and amino acid residues

To investigate Jacalin diversity in relation to the domain structure of TNJ, we annotated full-length protein sequences of representative *TNJ* genes with diverse Jacalin domain integrations in Geneious Prime v2022.1 (http://www.geneious.com) using the Pfam database (Mistry *et al*., 2020). Copies of the Jacalin domain were numbered according to their position in the protein for downstream analysis and interpretation of integration events.

To assess the conservation of key functional motifs, we aligned the NB-ARC domains of representative TNJ proteins from across the study species using MAFFT v7.487 (Katoh and Standley, 2013). We also extracted and aligned their corresponding TIR domains to examine whether critical residues known to mediate TIR-dependent defence signalling, including cysteine, glutamic acid, and tyrosine residues involved in the TIR catalytic activity (Huang *et al*., 2022; Jia *et al*., 2022; Lapin *et al*., 2022; Yu *et al*., 2022), are conserved in TNJ^TIR^ proteins. Additional NB-ARC and TIR domain proteins of L7, RPS4, RPP1 and RBA1 were included as references (Yu *et al*., 2022). The alignments were visualised and annotated using BioEdit v7.2.6.1 (Hall, 1999). To model TNJs and predict individual Jacalin domain structure, we employed AlphaFold3 model (Abramson *et al*., 2024) using RPP1:PDB 7CRC, one protomer from tetramer complex (Ma *et al*., 2020).

### Phylogenetic analysis of Jacalin containing NBS-encoding genes

To test whether TNJ proteins were monophyletic, we classified the 9,614 NBS-containing proteins annotated across eight species representing four major tribes (Thornhill et al., 2015; Balbinott et al., 2022) into *TNL, CNL, RNL*, and *TNJ*. The NB-ARC regions of 9,614 proteins were clustered into 4,911 non-redundant sequences using CD-HIT (-c 0.9 -n 5) (Fu *et al*., 2012). We aligned the non-redundant sequences with MAFFT v7.487 (Katoh and Standley, 2013), trimmed with ClipKIT (Steenwyk *et al*., 2020), and used maximum likelihood in IQ-TREE v2.0.3 (Minh *et al*., 2020) to infer the gene tree.

To determine whether Jacalin domains in *TNJ* genes arose from a single or multiple integration events, we aligned 418 integrated and non-integrated Jacalin domains annotated from the proteome and inferred their phylogenetic relationships. We extracted 522 protein sequences of individual Jacalin domains for analysis alongside 135 non-TIR-bound Jacalin domains (those lacking TIR and LRR regions) annotated across the eight species. All sequence alignments and phylogenetic analyses followed the procedures outlined above and were visualized using iTOL v4.8.1 (Letunic and Bork, 2024).

### Identification of hypervariable sites in TNJ domains

To assess sequence variability across TNJ domains, NB-ARC regions of 123 TNJ proteins from eight Myrtaceae species were aligned using MAFFT v7.487 (Katoh and Standley, 2013), and gappy columns were removed with ClipKIT (Steenwyk *et al*., 2020). Sequences were grouped into five clades using IQ-TREE-based maximum likelihood phylogeny. The full-length amino acid sequences were realigned within each group, and Shannon entropy scores were calculated (Stewart *et al*., 1997). Positions with entropy ≥ 1.5 were considered hypervariable (Prigozhin and Krasileva, 2021), and all groups contained such sites. To test whether hypervariable sites (HVS) were enriched in Jacalin domains, 10,000 random permutations were used to test for statistical enrichment within the TNJ alignment for each group.

### Analysis of site-specific positive selection

We tested for positive selection on each of the five TNJ phylogenetic groups used for HVS detection. The corresponding CDS of the full-length amino-acid sequences in each phylogenetic group were extracted with a custom Python script. We constructed CDS alignments from their corresponding aligned amino-acid sequences using pal2nal (Suyama *et al*., 2006), and gaps were visualized and removed using BioEdit v7.2.6.1 (Hall T, 1999). Sequences with < 60% identity were excluded to avoid alignment errors and incorrect estimates of evolutionary rates (Hodgins *et al*., 2016). Refined alignments were used to build maximum likelihood trees with IQ-TREE v2.0.3 (Minh *et al*., 2020). We utilized DAMBE7 (Xia, 2018) to assess Chi-square test for base frequency heterogeneity and substitution saturation by plotting transitions and transversions against Felsenstein (1984) F87 genetic distance to ensure alignment quality.

Because only a fraction of sites experiences positive selection over evolutionary time (Golding and Dean, 1998; Hodgins *et al*., 2016), pairwise comparisons averaging across all codons often miss such signals (Yang *et al*., 1998; Yang and Nielsen, 2000). To address this, we used a neighbour-joining tree based on the Goldman and Yang (1994) codon model to perform site-specific selection analysis (Yang and Swanson, 2002) and estimate dN/dS ratio with CODEML program of PAML (Yang *et al*., 1998; Yang and Nielsen, 2000). Equilibrium codon frequencies were estimated using F3X4 (Hodgins *et al*., 2016). Sites under positive selection (dN/dS > 1) were identified by likelihood ratio tests comparing M1a vs M2a and M7 vs M8 models, following Yang (1998) and Yang et al. (2005). To determine specific residue sites likely to have undergone diversifying selection, we used model M8, which supplements an adaptable null model provided by M7 beta model, with an additional site class of free *ω* ratio estimation (Yang and Swanson, 2002)

To assess the statistical significance of differences in positive selection across domains, we conducted likelihood ratio tests (LRT) on nested codon models using the χ² distribution at 0.05, 0.01, and 0.001 significance levels (Yang and Nielsen, 2000; Yang et al., 2005). Positively selected amino acid sites were identified using Bayes Empirical Bayes (BEB) inference following the recommendation in Yang *et al*. (2005). To ensure model consistency, analyses were repeated with fix_blength = 0 and varying starting ω values. Domain boundaries were defined via Pfam annotations, and mean ω with posterior probabilities was plotted across TIR-NBS-Jacalin domains using ggplot2 in R (v4.0.3; R Core Team, 2020).

## Results

### NBS-encoding gene and their NCD diversity in Myrtaceae

To gain insights into the variation of the NBS-encoding gene family among Myrtaceae species, NBS-encoding genes were annotated in the long-read-based genomes of eight representative species from four tribes of Eucalypteae, Myrteae, Zyzygieae and Melaleuceae (Table **S1**). Assembled genome sizes ranged from 270 Mb to 1073 Mb, with annotated protein counts ranging from 12,040 to 55,025 (Table **S4**). Of the total 226,152 genes predicted, 15,669 encoded NB-ARC domains, of which 11,657 (74.39%) contained both NBS and LRR domains (Table **S4**). The proportion of NBS-encoding genes ranged from 3.69% in *P. guajava* to 8.80% in *R. argentea*, representing an average of 6.89% of the predicted genes across the genome of eight species (Table **S5**; Fig. **S1**). *TNL*-type genes were the most abundant class detected in the Myrtaceae (Fig. **S1**), accounting for 50.0% of the full-length NBS-encoding genes (Table **S4**) and 2.78% of all annotated genes in the genome on average across the eight species (Table **S5**). The partial NBS-encoding genes were dominated by *N* and *TN* type (Table **S5**).

To determine the NCDs fused within NBS-encoding genes of each annotated species, we parsed the PfamScan outputs of the annotated gene models. We then identified the unique NCDs and their abundance for each species and NLR gene class. Of the genes annotated across the Myrtaceae species, many were found to encode for additional NCDs (Fig. **1a**). Across the eight Myrtaceae genomes annotated, we identified 181 unique NCDs (Table **S6**) occurring 1709 times in 810 (5.28%) of the NBS-containing proteins (Table **S4** & **S7**). While TNL proteins had the most diverse NCD complement, including 107 of the 181 unique NCDs identified (Fig. **1b**), the TN-type exhibited the greatest expansion of NCDs, occurring in 12% of TN proteins (199 of 1650; Fig. **1a**). This was largely due to the extreme expansion of the single dominant domain, Jacalin in TN and N-type genes (Fig. **1c**; Table **S6**). Of the 635 Jacalin copies detected, 522 were detected in 123 genes of TN-type, and 98 in 29 genes of N-type accounting for the largest proportion of the annotated Jacalins (Fig. **1d**).

**Fig. 1.**
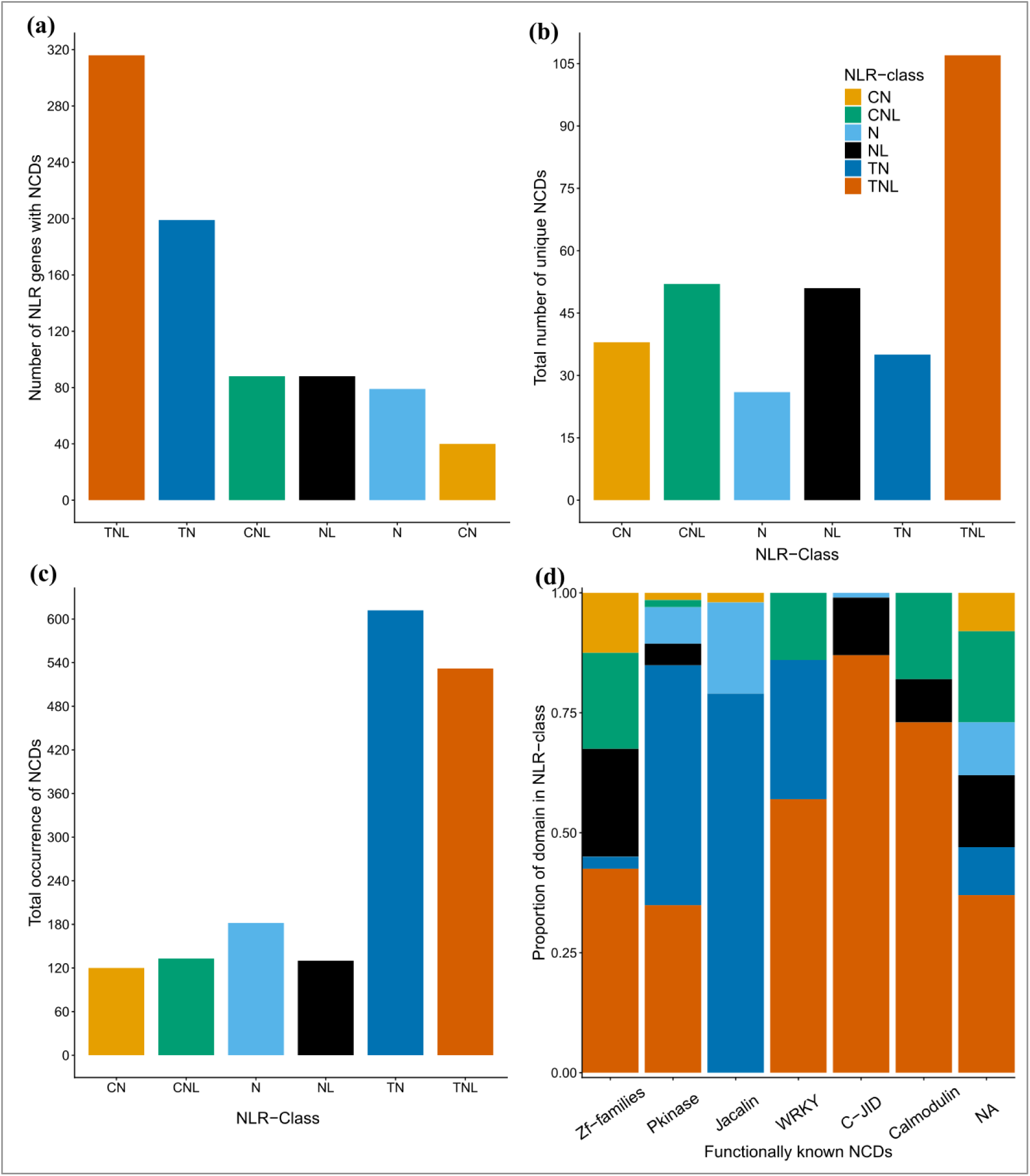
Expansion of NBS-encoding genes with non-canonical domains (NCD) in the genome of eight Myrtaceae species. **(a)** Percent of NCD containing NBS-encoding genes across the eight species in each *NLR* gene classes. **(b)** Total number of NCDs with unique Pfam identifiers in each NBS-encoding genes class. **(c)** Total number of occurrences of NCDs in each NBS-encoding gene class. **(d)** Proportion of occurrence of functionally known NCDs (zf-familiesy domains, Pkinase, Jacalin, WRKY, C-JID and Calmodulin-bind) in each NBS-encoding gene class. The definition of each NBS-encoding gene class is as described in Table **S3**.

In addition to the Jacalin domain, other functionally characterized protein domains, including Pkinase, C-JID, WRKY, Calmodulin-bind and zf-family domains, including the DNA binding zf-BED, were among the most abundant NCDs in NBS-encoding gene classes (Fig. **1d**; Table **S7**). While Pkinase and zf-family domains found in more diverse NBS-encoding genes, the annotated C-JID, Calmodulin-bind and WRKY tended to be found in TNL-type genes (Fig. **1d**).

### Jacalin domain structure and diversity in TNJ proteins

Analysis of TNJ domain structure and diversity revealed that multiple copies of Jacalin frequently occurred in the C-terminal of TIR-containing proteins that lacked LRR domains across the eight Myrtaceae species, predominantly in *Eucalyptus*, *Angophora* and *Syzygium* (Fig. **2a**). The Jacalin expansion varied greatly among Myrtaceae species, ranging from 13 copies in 3 genes in *P. guajava* to 121 copies in 30 genes in *E. albens* (Fig. **2a**; Table **S8**).

**Fig. 2.**
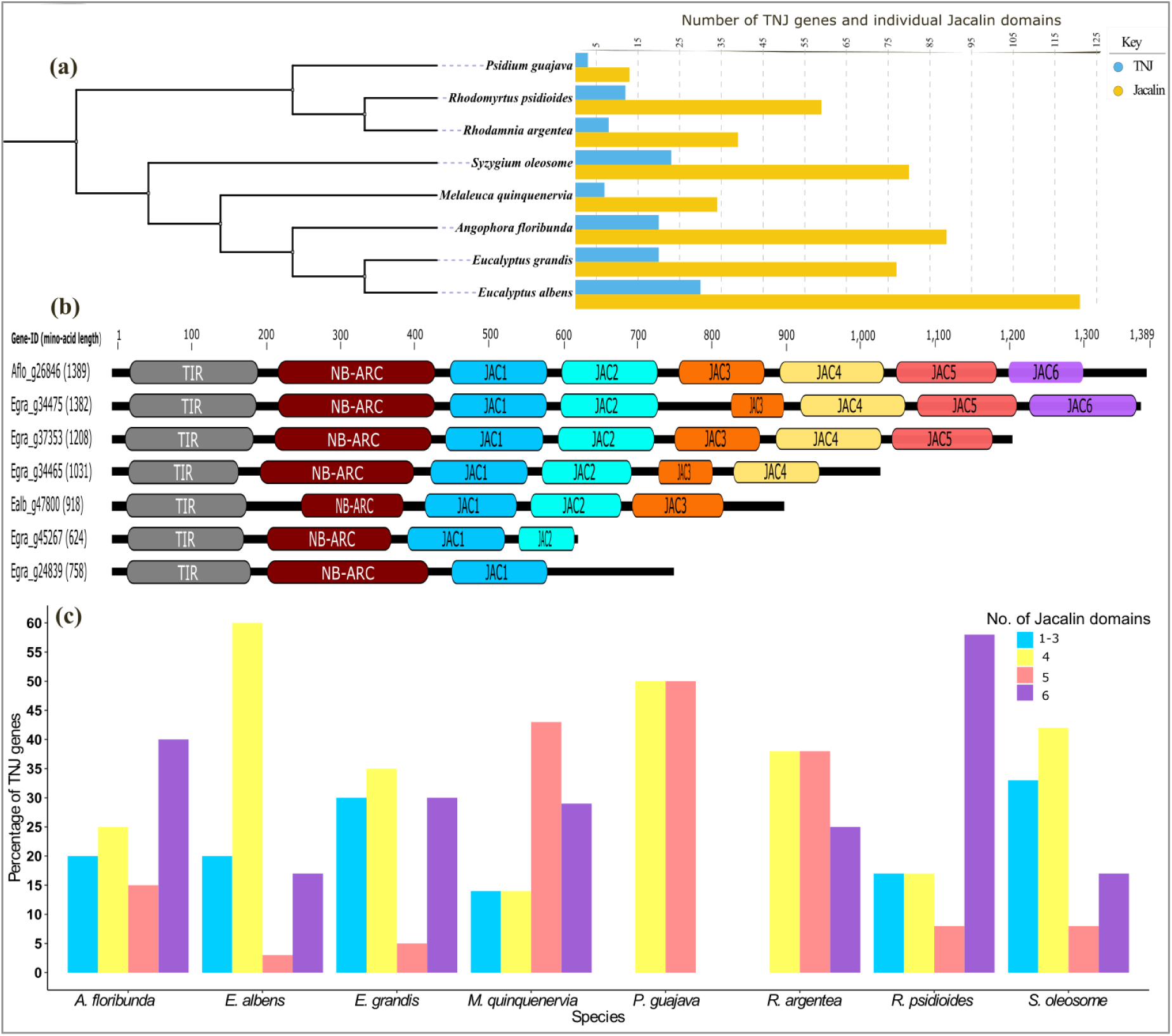
TNJ domain structure and diversity. **(a)** Taxonomy-based dendrogram of Myrtaceae species with their respective numbers of canonical *NLR* gene classes including *TNJ* and number of individual Jacalin domains annotated. **(b)** Representative structures of TNJ domains demonstrating integration of one–to-six individual Jacalin domains per gene. **(c)** Percentages of *TNJ* genes with different numbers of Jacalin domains in each genome. Genes with one and two Jacalin domains appeared with low frequency; and were grouped into those with three Jacalin domains for visualisation.

Sequence analysis suggests varied domain structure among *TNJ* genes, with between one and seven tandemly repeated Jacalin domains (Fig. **2b**), however, AF3 structural modelling revealed only six and that some Jacalin domains appeared to contain large insertions (Fig. **S2b**). Jacalin domains were characterised as 12 stranded, consisting of 3x4-stranded Beta-sheets. The sixth domain contained an unusually large loop/helix between the fifth-sixth and the nineth-tenth strands (Fig. **S2b**). Genes encoding one or two Jacalin domains were relatively rare as compared to those encoding with three, four, five or six Jacalins (Fig. **2c**). The average length of the Jacalin domain-containing region ranged from a minimum of 441 amino acids in *S. oleosome* to a maximum of 603 in *R. psidioides*, with an overall average of 539 across the species (Table **S8**). The average lengths of individual Jacalin domains varied from 102 amino acids for Jac3 to 131 for Jac4, although the amino-acid length varied within Jacalin positions (Fig. **2b**; Table **S9**).

### Functional motifs and amino-acid residues in TNJ

To investigate whether TNJ had retained the capacity to function as an *R* protein, we determined whether the functional motifs and amino-acid residues that have been reported to mediate the defence signalling response are conserved in its canonical domains. Inspection of the aligned amino-acid sequences of NB-ARC regions of the TNJ proteins showed that they possess the functionally important protein motifs, including the P-loop, Kinase-2, RNBS-B and GLPL motifs (Fig. **3a**). Conservation of functional amino-acid residues modulating TIR-mediated cell death was also confirmed in the amino acid sequence alignment of TIR regions of TNJ, following comparison with the reference sequences. The positions of important dimerization interfaces and catalytic amino-acid residues were identified in the alignments (Fig. **3b**). TNJ^TIR^ contained conserved residues predicted for both αA and αE (AE interface) and αD and αE (DE interface), which are required for TIR self-association and cell-death function of plant TIR domains (Zhang *et al*., 2017; Yu *et al*., 2022). Phenylalanine (F17) residue, which form a hydrogen-bonded interaction backbone and an element for stacking arrangements in the AE interface regions and TIR-domain self-association, was conserved in TNJ^TIR^ domains (Fig. **3b**). Also conserved in TNJ^TIR^ were the basic residues in DE interface region including lysine and arginine, and the cysteine residue (C73) in AE interface, vital in 2’3’-cAMP/cGMP synthetase activity, and glutamic acid residue (E76), required by TIR-domains to function as NADase to transduce recognition of pathogen into a cell death response (Fig. **3b**). Structural analysis of full-length TNJ protein shows that TNJ appears to be completely missing the ARC2/WHD subdomain that is associated with the NBS domains in TNL and CNL proteins (Fig. **S3**). ARC2/WHD subdomain is known to contain RNBS-D motif and the well described MHD domain.

**Fig. 3.**
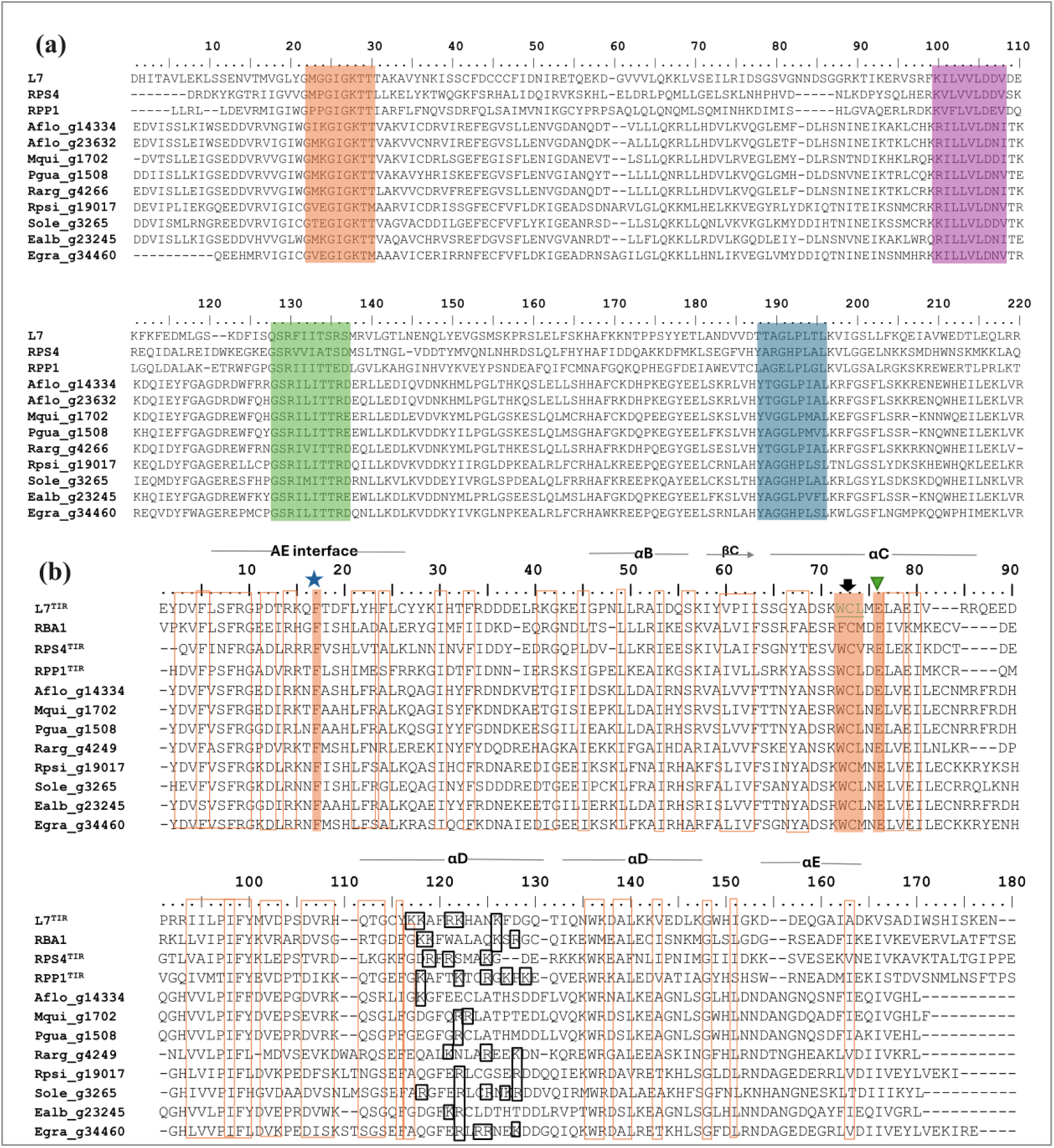
Sequence alignment of NB-ARC and TIR regions of representative TNJ proteins with reference proteins: L7, RPS4, RBA1 and RPP1 from NB-ARC of TNL proteins. **(a)** The conserved functional amino-acid motifs of the NB-ARC region are P-loop (orange), Kinase-2 (pink), RNBS-B (green), and GLPL (light blue). **(b)** Sequence alignment of TNJ^TIR^ domains showing the positions of TIR-self association AE and DE interfaces and conserved important amino-acid residues. Conserved residue at the AE interface are indicated by the blue star, and the basic residues in αD are indicated by black frames. The cysteine residue that is important in the 2’3’-cAMP/cGMP synthetase activity of TIR-mediated cell death (Yu et al., 2022) is conserved in TNJ^TIR^ and indicated by inverted black arrow. The glutamic acid residue that is required by TIR-NLR to function as NADase to transduce recognition of pathogen into a cell death response (Wan et al., 2019), is conserved in TNJ^TIR^ and indicated by inverted green triangle. All amino-acid sequences alignments in the Figure except L7^TIR^, RBA1 and RPS4^TIR^ were amino-acid sequence alignments of TNJ^TIR^. RBA1 is a TIR-only domain.

### Phylogenetic analysis of TNJ in the Myrtaceae species

To determine whether TNJ proteins form a monophyletic clade and infer the phylogenetic relationships between individual Jacalin domains, we constructed a phylogenetic tree of NB-ARC regions from *TNL, CNL, RNL* and *TNJ* genes, and individual Jacalin domains annotated from the eight Myrtaceae species (Fig. **4**). Phylogenetic analysis of NB-ARC regions revealed that *TNJ* genes form a monophyletic clade within the TNL-dominated clade (Fig. **4a**). Further phylogenetic grouping of NB-ARC regions of TNJ genes determined five major clades, which were used for downstream analysis of hypervariable sites and positive selection (Fig. **4b**).

**Fig. 4.**
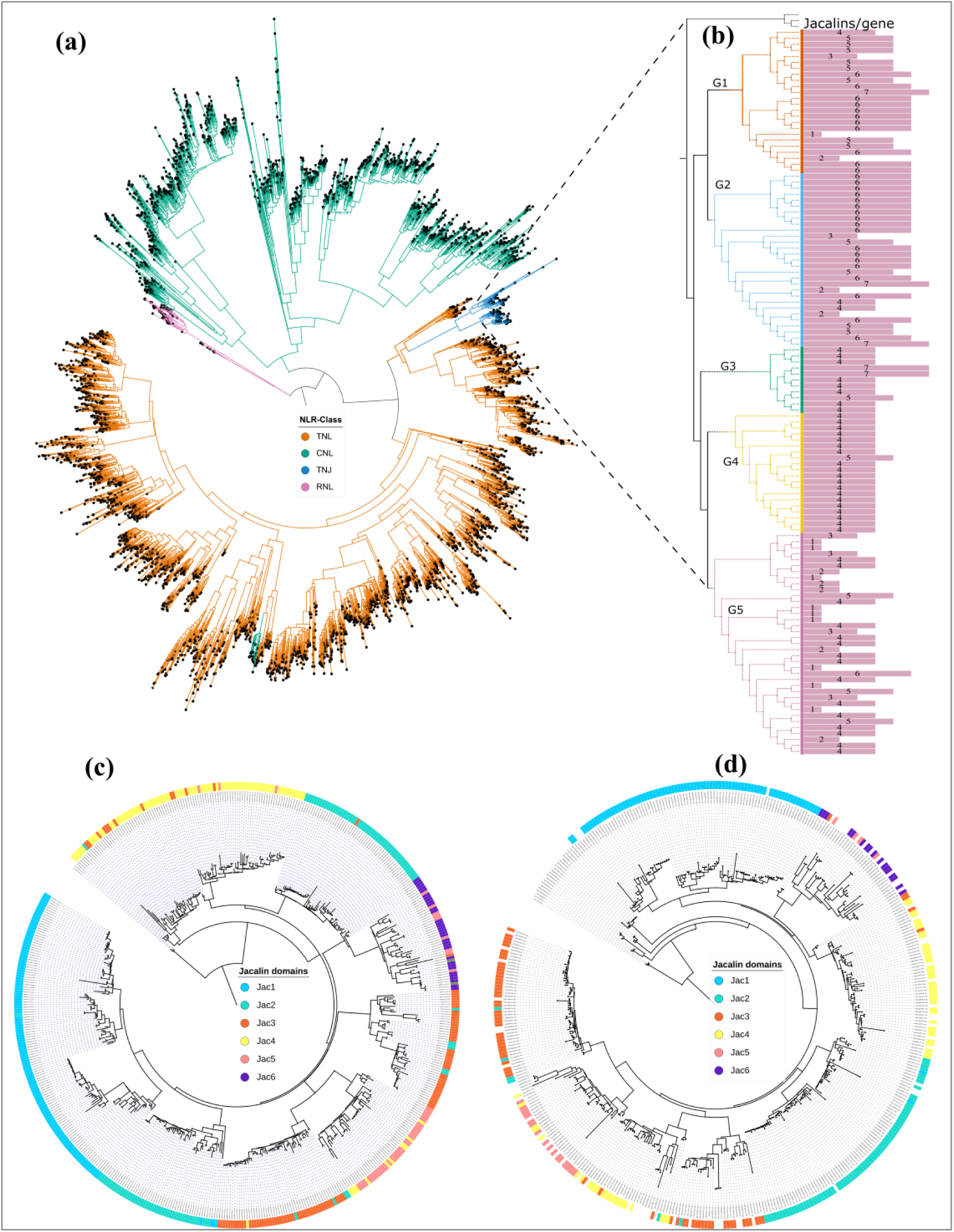
Phylogenetic trees of full-length *NLR* and *TNJ* proteins, and individual Jacalin domains from eight Myrtaceae species. **(a)** a maximum likelihood phylogenetic tree based on alignment of amino-acid sequences of NBS-ARC regions of 4911 proteins from *TNL, CNL, RNL* (outgroup) and *TNJ* gene classes from eight representative species of four major tribes across the Myrtaceae, including Syzygieae, Melaleuceae, Eucalypteae, and Myrteae. Branches are colour coded according to gene classes. **(b)** TNJ subtree shown with the corresponding numbers of individual Jacalin domains per gene, highlighting five phylogenetic groups (G1–5) used for downstream analysis. **(c)** A maximum likelihood phylogeny of individual Jacalin domains in TNJ proteins annotated from eight representative Myrtaceae species. **(d)** A maximum likelihood phylogeny of integrated (coloured bars) and non-integrated (white bars) Jacalin domains with Jacalin domains from *T. aestivum* as outgroups. The colour of the bars of each individual Jacalin domain around the tree were displayed by the colour of its corresponding positions in the protein sequence as indicated in Fig. 2b.

The phylogenetic relationships of individual Jacalin domains encoded in the *TNJ* genes shows that they tended to cluster by position in the gene, with one or two clades dominated by each position (Fig. **4c**). While the first (Jac1), fourth (Jac4), fifth (Jac5) and sixth (Jac6) positions tended to form a single clade each, the second (Jac2) and third (Jac3) positions formed two clades each (Fig. **4c**). In addition, less frequent Jacalin positions within clades tended to be produced by forward shifts (e.g., less frequent Jac4 domains in the Jac5-dominated clade). The phylogenetic relationships between individual Jacalin domains from non-TIR bound *N* genes of Myrtaceae tended to have similar grouping patterns as Jacalins within *TNJ* (Fig. **S4**).

Phylogenetic relationship of integrated and non-integrated Jacalin domains reveals the non-integrated Jacalin appeared to be ancestral (Fig. **4d**). The first (Jac1) and second (Jac2) integrated Jacalin domains appeared to form separate clade each, whereas from the third (Jac3) and later integrated Jacalin domains were not monophyletic, but instead were intermingled within the non-integrated Jacalin domains (Fig. **4d**). In addition, individual Jacalin domains from *CNL* genes of *O. sativa* also tended to form similar clustering pattern with Jacalin domains from TNJ (Fig. **S4**).

### Identification of hypervariable sites in Jacalin domain

To determine the distribution of hypervariable sites (HVS) among the TIR, NBS and Jacalin domains, domain boundaries in each TNJ phylogenetic group were estimated using their corresponding Pfam domain annotation (Table **S10**) and Shannon entropy analysis conducted. Of the 307 HVS detected across all domains in the phylogenetic groups, 251 (81.8%) were in the Jacalin region, and 43 (14.0%) and 13 (4.2%) were in the NB-ARC and TIR regions, respectively (Table **S11**). The total number of HVS and their distribution across domains varied among TNJ groups (Table **S11**). While G3 and G5 harboured few HVS on their Jacalin domains, they were concentrated in the Jacalin region in G1, G2 and G4. The Jacalin region was significantly enriched (p ≤ 0.001) for HVS for G1 and G2 (Fig. **5a**; Table **S11**).

**Fig. 5.**
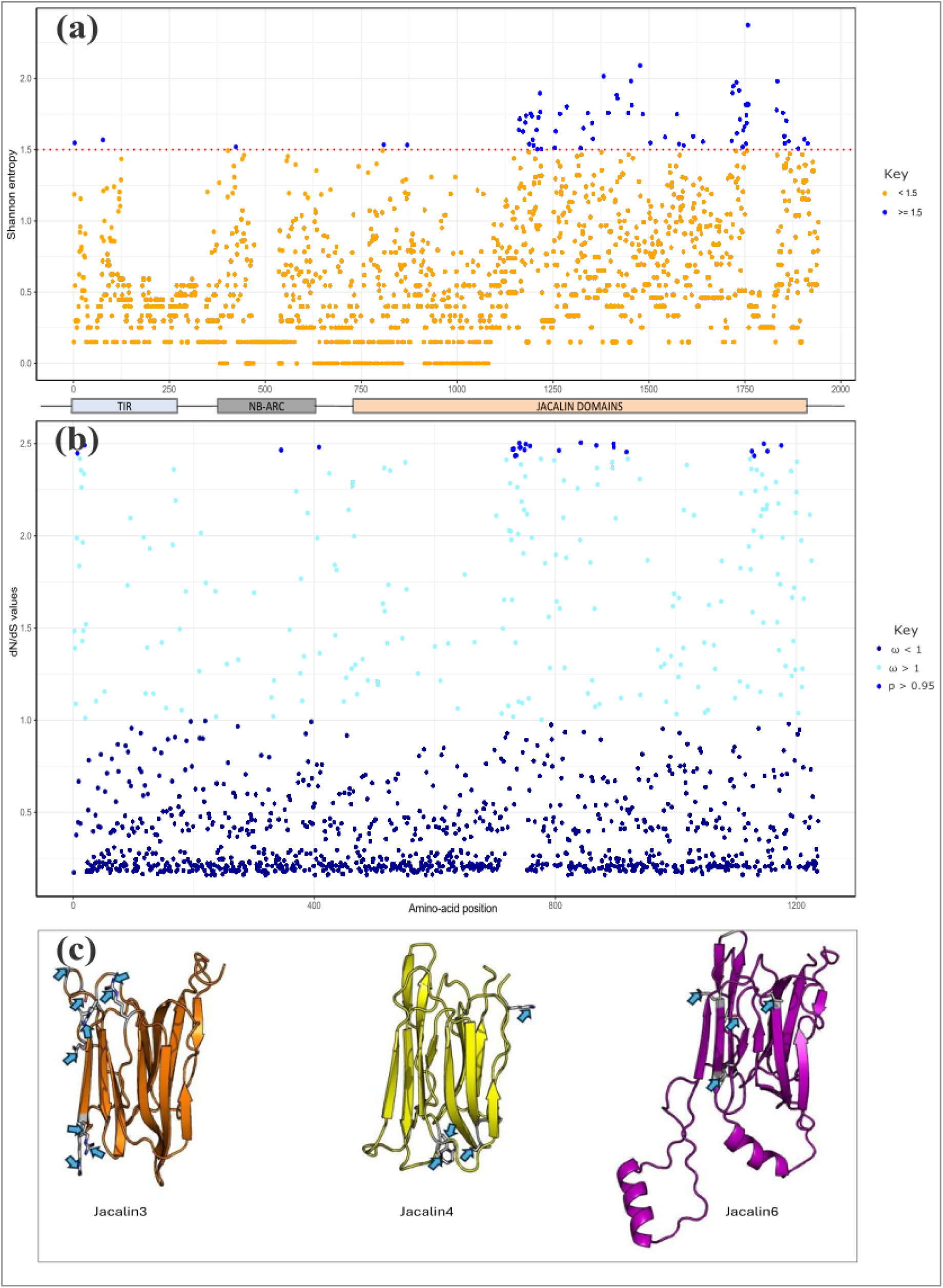
Shannon entropy and positive selection analysis of the group two monophyletic TNJ group. **(a)** Amino-acid residues with hyper-variable positions (entropy ≥ 1.5) were shown as blue points above red dotted line. **(b)** Mean dN/dS (ω) values and their corresponding posterior probabilities for positively selection (ω > 1) in TNJ proteins. Amino-acid positions with posterior probability > 0.95 were likely undergone significant positive selection and are indicated by blue points in the plot. Positively selected sites were detected by comparing M7 and M8 site models in CODEML. **(c)** AlphaFold 3 structural model analysis result showing examples of Jacalin 3, 4 and 6 domains that underwent positive selection were surface-exposed. Surface-exposed sites were indicated by light blue arrows. Domain boundaries were estimated from Pfam domain annotation and indicated using boxes under the plot.

### Detection of positive selection in TNJ

To evaluate the distribution of sites showing signatures of positive selection, genes in each *TNJ* group were subjected to evolutionary model comparisons. Transition (t) and transversion (v) plots and Chi-square tests for heterogeneity suggested that the five phylogenetic groups of TNJ sequences were suitable for positive selection analysis. Nucleotide frequencies did not differ significantly among sequences in any phylogenetic group of *TNJ* genes (all *p* > 0.97; Table **S12**,), and transition/transversion ratios did not indicate saturation of substitutions (Fig. **S5**).

Of the five groups analysed, four had significant likelihood ratio tests (LRT) under comparison of models M1a (neutral) and M2a (positive selection) (Table **S13**). Sequences in G5 had no residue positions with *ω* significantly elevated, suggesting that purifying selection had predominated in that group. The proportion of sites with signatures of positive selection identified by comparison of M8 and M7a ranged from 0.11% in G3 to 2.35% in G2, followed by 1.85% in G4 (Table **S14**).

The majority (76.5%) of sites detected with positive selection signature were in the Jacalin region of the TNJ, followed by 13.7% and 9.8% in TIR and NBS regions of the domains, respectively (Table **S13**; Table **S14**; Fig. **5b**; Fig. **S6**).

To determine whether the amino acid residues associated with positive selection in phylogenetic group G2 of TNJ are surface-exposed, we employed AlphaFold3 structural analysis. Our results reveal that the majority (∼80%; 21/26) of sites with signatures of positive selection are located within the Jacalin region, with 10 in Jacalin 3, 6 in Jacalin 4, and 5 in Jacalin 6. Among these, 20 residues are surface-exposed (Fig. **5c**).

## Discussion

### NBS-encoding genes are abundant in Myrtaceae

The frequency and severity of disease outbreaks in forest trees have intensified in recent years, driven by range expansions, the introduction of new species, and infections in previously unaffected hosts becomes more common, often resulting in severe damage or even tree mortality (Tobias *et al*., 2016; Teshome *et al*., 2020). Against this backdrop, our study examines the significance of a previously uncharacterised NLR architecture in Myrtaceae, defined by the integration and expansion of a Jacalin non-canonical domain at the C-terminal of TIR–NBS proteins (TNJ). By examining the evolutionary and functional implications of the novel TNJ proteins, this study provides insight into how Myrtaceae species may have diversified their NBS-encoding protein repertoires in response to emerging threats.

The Myrtaceae genomes examined here have relatively high numbers of predicted *NLR* genes, with an average of 2625 NB-ARC domains; representing 6.29% of all predicted genes among the eight species. This substantially surpasses counts in most other angiosperms (Yang *et al*., 2008; Shao *et al*., 2014; Jia *et al*., 2015; Sarris *et al*., 2016; Shao *et al*., 2016; Baggs *et al*., 2017; Van de Weyer *et al*., 2019; Xue *et al*., 2020; Liu *et al*., 2021b). The high percentage (6.29%) of predicted NBS-encoding genes demonstrate the importance of long-read assembled genomes for resistance gene analyses when compared to earlier findings of ∼1% within short-read assembled genomes (Tobias and Guest, 2014). This observation aligns with findings that many long-generation tree species, including *Quercus rubra* and *Q. lobata*, exhibit higher proportions of genes encoding the NB-ARC domain than many other plant species (Ngou *et al*., 2022).

The Myrtaceae species examined have undergone lineage-specific expansion of the *TNL*-type gene classes. The *TNL*s make up 50.0% of the full-length *NLR* genes across the species studied here, with the rest being *CNL, RNL* and *NL* class genes. In contrast, a lower TNLs than CNLs has been reported in *Vitis vinifera* (12.4%) and in species of the Fabaceae (30.5 - 38.7%) and Solanaceae (5.2% - 15.2%) families (Shao *et al*., 2016). *NLR* gene classes have undergone lineage-specific expansion and contraction in other groups (Liu *et al*., 2021b). For instance, *RNLs* have expanded independently in Rosales and Conifers (Jia *et al*., 2015; Van Ghelder *et al*., 2019), while *CNLs* have expanded more than *TNLs* in 268 of the 305 angiosperm genomes (Liu *et al*., 2021a) and in various Solanaceae species (Jupe *et al*., 2012; Seo *et al*., 2016; Seong *et al*., 2020; Guo *et al*., 2025). Multiple lineage-specific expansions of NLR subgroups, suggest that common forces, including gene duplication and diversifying selection are driving NLR diversification across diverse plant lineages.

### The Jacalin domain is highly expanded in TIR-encoding *R* genes of Myrtaceae

NLR protein domain expansion is an evolutionary mechanism by which plants optimise their response to rapidly evolving pathogen (Song *et al*., 2017; Zhang *et al*., 2022; Wu and Derevnina, 2023). For instance, the LRR domains of *NLRs* have undergone diversifying selection on their solvent-exposed residues to maintain the ability to bind evolving effector proteins (Prigozhin and Krasileva, 2021). In this study, we show that Jacalin domains are highly expanded at the C-terminus of *TNJ* genes, which lack the typical LRR domains of full-length NLRs. Similar to the LRR domains of TNLs and CNLs, Jacalin shows repeated integration into TN genes, suggesting that Jacalin may have evolved as a potential functional analogue for the LRR in Myrtaceae. Phylogenetic analysis showed that Jacalin containing TNJ proteins formed a single clade, indicating a single ancestral Jacalin insertion into a *TNJ* gene, followed by possible subsequent tandem duplications. Consistent with this, NBS-encoding genes from bacteria, archaea, glaucophyte, algae, and bryophyte lineages, which were integrated with various repeats and kinase domains were clustered separately on the phylogenetic tree (Andolfo *et al*., 2019).

Jacalin-like lectin domains are implicated in responses to biotic and abiotic stresses, including defence (Song *et al*., 2014; Weidenbach *et al*., 2016; Esch and Schaffrath, 2017; Han *et al*., 2018; Ma *et al*., 2021). For instance, by obstructing the over-accumulation of viral RNA, a Jacalin containing protein from the *Arabidopsis*, JAX1, confers resistance against potexvirus (Yamaji *et al*., 2012). Recent studies reveal functional divergence of the carbohydrate-binding jacalin-like lectin *OsJAC1* depending on rice genetic background. In *O. sativa* ZH10 background, its overexpression confers broad-spectrum resistance by directing defence proteins to infection sites and enhancing immune responses (Weidenbach *et al*., 2016). In contrast, CRISPR/Cas9-mediated disruption of OsJAC1 in the *O. sativa* ZH11 background results in increased disease resistance (He *et al*., 2025), suggesting that its role in plant immunity is strongly influenced by genetic context and may involve complex regulatory interactions within distinct genomic backgrounds.

It is plausible that the Jacalin domains in TNJ proteins may substitute for LRRs in effector recognition. LRR domains are known for their high degree of recognition specificity and adaptability in binding various pathogen effectors via their hypervariable, surface-exposed amino acid residues (McHale *et al*., 2006; Prigozhin and Krasileva, 2021). In soybean, expansion of LRR has been suggested to contribute to broad-spectrum resistance by *Rps11* gene, a cluster of large *NLR* genes of a single origin, against *Phytophthora* root and stem rot caused by *Phytophthora sojae* (Wang *et al*., 2021). In *Arabidopsis thaliana*, recognition of glycan-based peptidoglycan (PGN) molecules by the lysin motif (LysM) receptors LYM1, LYM3, and LysM receptor kinase CERK1 triggers canonical immune responses and defence gene activation (Willmann *et al*., 2011). Given their high copy numbers, Jacalin domains may similarly facilitate resistance to diverse pathogens through direct effector binding as has been shown for LRRs, particularly through perception of molecular signatures such as glycans, a carbohydrate molecule shared across whole classes of microbial organisms (Wanke *et al*., 2021). In addition, in Myrtaceae, we confirmed up to six Jacalin copies per gene and as many as 123 Jacalin copies in a single genome (*E. albens*), suggesting a role in NBS-encoding protein diversification and adaptation to fast-evolving pathogens (Kroj *et al*., 2016; Sarris *et al*., 2016; Grund *et al*., 2019; Van de Weyer *et al*., 2019). Our study revealed that non-integrated Jacalin domains represent an ancestral lineage to the integrated forms with frequent escape and re-integration resulting in multiple integrated clades. This suggesting that Jacalin domains originally existed as standalone proteins and were later recruited into NLRs, however the domain integration is not terminal state but part of an ongoing process of birth-and-death evolution (Kroj *et al*., 2016; Zeng *et al*., 2025). This is consistent with the integrated decoy model, whereby host proteins targeted by pathogen effectors are duplicated and subsequently fused into NLRs, generating novel immune receptors (Cesari *et al*., 2014; Kroj *et al*., 2016; Contreras *et al*., 2023).

### Essential functional motifs and residues are conserved in TNJ proteins

The presence of conserved functional elements in TNJ suggests that the gene family has retained the capacity to function as R proteins. Four conserved motifs essential for ATP hydrolysis (Shao *et al*., 2016; Van Ghelder *et al*., 2019) are conserved in the NBS region of TNJ. P-loop, Pkinase-2, RNBS-B and GLPL motifs mediate nucleotide binding and transformation of the terminal phosphate group in the modern NTPase enzymes, causing signal activation and receptor oligomerization that finally culminates in the hypersensitive cell-death response and host immunity (Bork and Koonin, 1994; Danot *et al*., 2009; Shao *et al*., 2016; Romero Romero *et al*., 2018; Van Ghelder *et al*., 2019). Similar conservation was reported in angiosperm NLRs across Fabaceae, Brassicaceae, Solanaceae, and Poaceae, indicating that it should also be expected in any novel NLR subclass (Shao *et al*., 2016). The P-loop is of particular significancy, as it represents one of the earliest known nucleotide-binding signatures in eukaryotes (Zheng *et al*., 2016; Romero Romero *et al*., 2018). TNJs lack the RNBS-D motif, which appears to be poorly conserved even among CNL and TNL genes across 22 angiosperm genomes (Shao *et al*., 2016). In canonical TIR-CC-NLRs, the ARC2/WHD subdomain within NB-ARC region contains a well characterized MHD motif (Wang *et al*., 2015). Notably, mutation of the conserved histidine in the MHD motif often results in auto-immune response (van Ooijen *et al*., 2008; Wang *et al*., 2015). However, structural analysis of TNJs indicate that the ARC2/WHD subdomain is entirely absent. This structural loses suggests that TNJs are unlikely to be regulated through the canonical MHD-dependent molecular switch and instead may employ a distinct activation and regulatory mechanisms compared to the canonical TIR-CC-NLRs.

As is the case in TNL proteins generally, the TIR domain of *TNJ* genes in Myrtaceae shows high conservation of the catalytic glutamic acid (E), which is essential for TIR-based immune signalling. Upon pathogen recognition by the TNL protein, this residue catalyses the degradation of oxidized NAD⁺, initiating TIR-dependent cell-death signalling upstream of EDS1 and NRG1 (Horsefield *et al*., 2019; Wan *et al*., 2019). For example, purified TIR domains from canonical TNLs like L6 (flax) and RUN1 (grapevine) have demonstrated E-dependent cleavage of NAD⁺/NADP⁺ (Wan *et al*., 2019), resulting in the accumulation of TIR-based key signalling molecule, variant cyclic ADP ribose (v-cADPR) (Chang *et al*., 2019; Wan *et al*., 2019). Mutation of this conserved glutamic acid in proteins like RBA1 and BdTIR TIR-only domains, and in RPP1, L6, RUN1, SNC1, and RPS4 of TNL^TIR^ domains significantly impairs the cell-death phenotype in *Nicotiana benthamiana*, confirming the importance of NAD⁺ cleavage for TIR signalling (Yu *et al*., 2022). This suggests that TNJ proteins in Myrtaceae may also trigger immune signalling through TIR-mediated NADase activity. Furthermore, TNJ^TIR^ domains conserve key basic residues at AE and DE interface regions essential for TIR self-association and 2’3’-cAMP/cGMP synthetase activity (Zhang *et al*., 2017; Wan *et al*., 2019; Yu *et al*., 2022). These structural features, also found in other functional TNL^TIR^ proteins (Zhang *et al*., 2017; Chang *et al*., 2019; Wan *et al*., 2019; Yu *et al*., 2022), underline the potential of TNJ genes to mediate immune responses through conserved self-association interfaces, NADase enzymatic function, and induction of cell death signalling in response to pathogen challenge.

### TNJ contain hyper-variable and positively selected residues clustered in the jacalin domain

In this study, the sequence diversity of phylogeny-based TNJ orthologous groups was assessed using Shannon entropy and codon-specific tests for positive selection (Stewart *et al*., 1997; Yang, 1998; Yang *et al*., 2005; Prigozhin and Krasileva, 2021). This helped to identify TNJ clades with more HVS and harboured residues with higher dN/dS ratios than others, suggesting that there could be a clade-specific effector recognition specificities evolving under diversifying selection in Myrtaceae species. This contrasts with most NLRs, which exhibit conserved functions and limited sequence diversity in entropy-based analyses, including the indirect effector recognizer CNL RPS2, the conserved helper RNL NRG1.1, and the TNL RRS1B, which employs an integrated WRKY domain (Prigozhin and Krasileva, 2021).

NBS-encoding receptor clades with more HVS are proposed to evolve more rapidly (undergo diversifying selection) than those clades with less HVS (Krasileva *et al*., 2010; Prigozhin and Krasileva, 2021), and they have been associated with previously characterised known proteins of direct binders. For instance, 16 *TNL* genes from the hypervariable *RPP1* clade and 14 *CNL* genes from the *RPP13* clade in *Arabidopsis thaliana* revealed periodic entropy peaks in their LRR region, a molecular signature that aligns with their function in direct, rather than indirect recognition of effector proteins (Prigozhin and Krasileva, 2021). In addition, LRR domains of 11 sub-groups of LRR-RLK proteins have been found to evolve more rapidly than their cognate RLK domains (Man *et al*., 2023). Consistent with this, the distribution of positively selected residues was predominantly clustered on the predicted Jacalin domain of TNJs, suggesting Jacalin domains have evolved with signatures of selection similar to what has been observed in LRR domains in other species (Prigozhin and Krasileva, 2021; Man *et al*., 2023) and potentially provide similar pathogen recognition specificity function. Hence, Jacalin and LRR are potentially functional analogues in terms of pathogen molecule recognition.

In addition, the predominantly surface-exposed localization of the positively selected residues highlights the presence of solvent-accessible and highly diversified interfaces. These interfaces are likely involved in mediating ligand binding (Ma *et al*., 2003). These key pieces of evidence suggest that *TNJ* could potentially be a new class of *R* genes that has evolved in Myrtaceae family with distinct characteristics alongside conserved functional motifs and amino-acid residues. Consistent with this, new *R* gene classes with a distinct structure have been reported in recent years. These include the NL-only domains (Van de Weyer *et al*., 2019), the autonomous and ancient *NLR* (ANL) that encode an auto-active coiled-coil (CC) domain (Lee *et al*., 2020), and the zf-BED-NBS-encoding novel resistance profile that was found to form a distinct phylogenetic clade in poplar (Yang *et al*., 2008).

The distinct clustering of first and second integrated Jacalin domains into separate clades suggests the early and independent integration event of Jacalin domains into the NLR proteins with subsequent lineage-specific duplication and divergence. Contrary to this, the nesting of the third and later integrated Jacalin domains among non-integrated Jacalins indicates that the subsequent integrations were more recurrent, possibly involving ongoing domain shuffling and recruitment events from the proteome, contributing to the modular evolution of Myrtaceae NBS-encoding genes. This is consistent with the hypothesis that a group of Jacalin domains became integrated into a single ancestral TN domain and expanded from one point, with multiple integrated copies likely arising from tandem duplication. In contrast, Marchal *et al*. (2018) showed substantial divergence between integrated and non-integrated zf-BED domains of *T. aestivum* suggesting that integrated BED domain had potentially evolved as a result of multiple integration events. Similar to their phylogenetic relationship in Myrtaceae, integrated Jacalin domains in CNL genes of *Oryza sativa* appear to follow grouping patterns according to their position in the protein. This common trend suggests either a shared ancestral Jacalin integration before the monocot-eudicot divergence ∼140–150 Mya (Johnson and Thomas, 2007), followed by lineage-specific expansion into CNL and TN type proteins or convergent evolution via independent integration events in Poaceae and Myrtaceae lineages.

In conclusion, our study reveals that Jacalin domain, notably abundant in Myrtaceae family, occurs multiple times per gene at the C-terminal of TIR-type NBS-encoding genes. Domain and phylogenetic analyses reveal that TNJ genes form a distinct clade, and individual Jacalin domains group according to their position within the tandem repeats, suggesting sequential expansion after the initial integration. In most phylogenetic groups, Jacalin is highly polymorphic, likely enabling rapid evolution of novel binding specificities to match fast-evolving effectors. Despite the absence of the ARC2/WHD motif, the conservation of other functional motifs and the presence of surface-exposed positively selected residues within Jacalin domain supports its functional analogy to LRR domains, possibly mediating pathogen recognition or oligomerization. These findings imply that different plant families may evolve distinct recognition domains and clade-specific *R-*gene architectures contributing to niche adaptation. Future functional studies on these genes and their regulatory networks should provide further elucidation of these aspects of the immune systems.

## Acknowledgements

The authors acknowledge Norman Gaywood of UNE and Peter Minogue of ANU, Research School of Biology, for facilitating access the computing facilities of the respective Universities.

## Author contribution

Conceptualization, methodology and investigation: TT, RA, BS, ShW; Data curation, NLR Analysis and Writing: TT, BS, RA, SW; Data visualization: TT, RA, XZ, SW; Supervision, reviewing and editing: RA, BS, ShW; Sampling, DNA extraction, ONT sequencing and genome assembly: AJ, SF, JB; NLR pipeline conceptualization and development: TT, BS, ZL, PT

## Data Availability

The resistance gene annotation tool is available at https://github.com/ZhenyanLuo/FindPlantNLRs and is registered on bio.tools (https://bio.tools/findplantnlrs). The interactive tree files can be viewed at https://itol.embl.de/shared/1cPvaR2fQsnjG.

## Supporting information

| File name | Description |
| --- | --- |
| ***Supplementary files.xlsx*** | **Fig. S1.** The proportion of each NLR class genes as expressed in percentage of the total predicted gene models in the genome of the eight annotated broader Myrtaceae species.<br>**Fig. S2:** Full length TNJ structure model demonstrating TIR and NBS domains, and individual Jacalin domains.<br>**Fig. S3:** AlphaFold3 structure of two full-length TNJ proteins from *Angophora floribunda* and *Eucalyptus grandis* compared to the known RPP1 protein.<br>**Fig. S4.** A maximum likelihood tree that shows phylogenetic relationship of non-TIR-bound integrated Jacalin domains (Jacalin domains from N proteins, which have no N-terminal and C-terminal domain) across the eight species.<br>**Fig. S5.** Plot of transition (t) and transversion (v) against the F84 distance for TNJ phylogenetic groups with sites of positive selection.<br>**Fig. S6.** Shannon entropy and positive selection analysis of TNJ phylogenetic groups.<br>**Table S1.** List of Taxonomic and GeneBank ID of the Myrtaceae species used for NBS-encoding genes annotation.<br>**Table S2.** BUSCO completeness of the genome assembly of the non-Eucaluptea species annotated. For assembly completeness of the Eucalyptea species in the study, refer to Ferguson et al., 2024).<br>**Table S3.** Definitions of NLR protein classes annotated using the FindPlantNLRs annotation pipeline<br>**Table S4.** Genome size, gene model predicted and NBS-encoding genes annotated in the genomes of eight representative species from Myrtaceae<br>**Table S5.** Proportion of centrally conserved NBARC, and full-length and partial type NLR genes in the genomes of eight Myrtaceae species.<br>**Table S6.** List of 181 unique non-canonical domains with their count of frequency of integration into the NLR genes of eight Myrtaceae species annotated.<br>**Table S7.** Complete list of integrated domains along with their PfamID and NLR classes identified in the eight Myrtacea species annotated.<br>**Table S8.** Table S8. Average start and end, and amino-acid sequence length of the Jacalin domains in the eight Myrtaceae species annotated.<br>**Table S9.** Average start and end, and overall mean amino-acid sequence length of individual Jacalin domain positions in the studied Myrtaceae species.<br>**Table S10.** TIR-NBS-Jacalin domain boundaries and mean sequence length in the alignment of each phylogenetic groups of TNJ estimated from their corresponding Pfam domain annotation.<br>**Table S11.** Proportion of amino acid residue with hyper variable sites in the TIR-NBS-Jacalin domains of TNJ proteins.<br>**Table S12.** Chi-square test of heterogeneity of nucleotide frequencies among the sequences using Yates correlation.<br>**Table S13.** Likelihood ratio test (LRT) and parameter estimate for five phylogenetic groups of TNJ sequences used for positive selection analysis using site-model of the CODEML program of PAML.<br>**Table S14.** Proportion of positively selected amino acid residue sites in TIR-NBS-Jacalin domains of TNJ. |

